## Supplementary file with additional data. for "Xanthomonas citri subsp. citri type III effector PthA4 directs the dynamical expression of a putative citrus carbohydrate-binding protein gene for canker formation"

Supplementary file 1. Supplementary file with additional data.

a. List of bacterial strains and plasmids used in this study

| **Strains or plasmids** | **Relevant characteristics** | **Resources** |
| --- | --- | --- |
| **Strains** | | |
| *Xanthomonas citri* subsp. *citri* | | |
| *Xcc* 29-1 | Wild-type strain isolated from *Citrus* *sinensis* in Jiangxi Province, China | This lab |
| *Xcc* 29-1/avrXa7 | Km^r^, *Xcc* 29-1 carrying pUFR034:avrXa7 | Sun *et al*., 2018 |
| Mxac 126-80 | Km^r^, a Tn5 insertion mutant of *pthA4* gene derived from *Xcc* 29-1 | Song *et al*., 2015 |
| *Xcc* 049 | Wild-type strain isolated from *C.* *sinensis* in Chongqing Province, China | This lab |
| *Xcc* 049E | A tale free strain of *Xcc* 049 | Ge *et al*., 2019 |
| *Xcc* 049E/pthA4 | Km^r^, *Xcc* 049E containing the *pthA* gene | Ge *et al*., 2019 |
| *Escherichia coli* | | |
| DH5α | *F^-^ recA hsdR17 (rk^−^, mk^+^) ϕ80lacZ∆M15* | Clontech |
| BL21(DE3) | *F^-^, ompT, hsdSB (rB^-^mB^-^), gal, dcm* | Novagen |
| Saccharomyces cerevisiae | | |
| EGY48 | *MATα, ura3, his3, trp1*, LexAop-LEU2 | Clontech |
| AH109 | *MATα, trp1*, *leu2,* lacZ, HIS3, ADE2, MEL1 | Clontech |
| *Agrobacterium tumefaciens* | | |
| GV3101 | Rif^r^, with Ti plasmid pMP90 | Koncz and Schell, 1996 |
| **Plasmids** | | |
| pHB | Km^r^, a binary vector to express gene under control of a double CaMV 35S promoter | Mao *et al*., 2005 |
| pHB-Cs9g12620 | Km^r^, the 1122-bp full-length *Cs9g12620* gene cloned in pHB | This study |
| pHB-CsLOB1 | Km^r^, the 714-bp full-length *CsLOB1* gene cloned in pHB | This study |
| pHB-PthA4 | Km^r^, the 3492-bp full-length *pthA4* gene cloned in pHB | This lab |
| pGD3FLAG-PthA4 | Km^r^, the 3492-bp full-length *pthA4* gene cloned in pGD3FLAG | This lab |
| pGWB435-P*Cs9g12620* | SH^r^, a 463-bp DNA fragment of *Cs9g12620* gene promoter region cloned in pGWB435-LUC | This study |
| pGWB435-P*Cs9g12620*-M | SH^r^, a 397-bp DNA fragment of *Cs9g12620* gene promoter region cloned in pGWB435-LUC | This study |
| pGWB435-P*Cs9g12620*-M_A_ | SH^r^, a 463-bp DNA fragment of *Cs9g12620* gene promoter region cloned in pGWB435-LUC | This study |
| pGWB435-P*Cs9g12620*-M_C_ | SH^r^, a 463-bp DNA fragment of *Cs9g12620* gene promoter region cloned in pGWB435-LUC | This study |
| pGWB435-P*Cs9g12620*-M_G_ | SH^r^, a 463-bp DNA fragment of *Cs9g12620* gene promoter region cloned in pGWB435-LUC | This study |
| pGWB435-P*Cs9g12620*-M_LB1_ | SH^r^, a 459-bp DNA fragment of *Cs9g12620* gene promoter region cloned in pGWB435-LUC | This study |
| pGWB435-P*Cs9g12620*-M_LB2_ | SH^r^, a 459-bp DNA fragment of *Cs9g12620* gene promoter region cloned in pGWB435-LUC | This study |
| pGWB435-P*Cs9g12620*-M_LB1/2_ | SH^r^, a 455-bp DNA fragment of *Cs9g12620* gene promoter region cloned in pGWB435-LUC | This study |
| pGWB435-P*Cs9g12650* | SH^r^, a 460-bp DNA fragment of *Cs9g12650* gene promoter region cloned in pGWB435-LUC | This study |
| pG221 | Ap^r^, Yeast one-hybrid plasmid | Ye *et al*., 2004 |
| pG221-P*Cs9g12620* | Ap^r^, a 463-bp DNA fragment of *Cs9g12620* gene promoter region cloned in pG221 | This study |
| pG221-P*Cs9g12620*-M_A_ | Ap^r^, a 463-bp DNA fragment of *Cs9g12620* gene promoter region cloned in pG221 | This study |
| pG221-P*Cs9g12620*-M_C_ | Ap^r^, a 463-bp DNA fragment of *Cs9g12620*gene promoter region cloned in pG221 | This study |
| pG221-P*Cs9g12620*-M_G_ | Ap^r^, a 463-bp DNA fragment of *Cs9g12620* gene promoter region cloned in pG221 | This study |
| pG221-P*Cs9g12620*-M_LB1_ | Ap^r^, a 459-bp DNA fragment of *Cs9g12620* gene promoter region cloned in pG221 | This study |
| pG221-P*Cs9g12620*-M_LB2_ | Ap^r^, a 459-bp DNA fragment of *Cs9g12620* gene promoter region cloned in pG221 | This study |
| pG221-P*Cs9g12620-*M_LB1/2_ | Ap^r^, a 455-bp DNA fragment of *Cs9g12620* gene promoter region cloned in pG221 | This study |
| pGADT7 | Ap^r^, GAL4(768–881) PADH1, TADH1, HA tag | Clontech |
| pGBKT7 | Km^r^, GAL4(1–147) DNA-BD, TRP1, c-Myc epitope tag | Clontech |
| pGADT7-pthA4 | Ap^r^, the 3492-bp full-length *pthA4* gene cloned in pGADT7 | This lab |
| pGBKT7-pthA4 | Km^r^, the 3492-bp full-length *pthA4* gene cloned in pGBKT7 | This study |
| pGBKT7-CsLOB1 | Km^r^, the 714-bp full-length *CsLOB1* gene cloned in pGBKT7 | This study |
| pCAMBIA1381 | Km^r^, plant binary expression vector carrying a promoterless *gusA* gene with start codon | CAMBIA |
| P*Cs9g12620*-GUS | Km^r^, a 463-bp DNA fragment of *Cs9g12620* gene promoter region cloned in pCAMBIA1381 | This study |
| P*CsLOB1*-GUS | Km^r^, a 584-bp DNA fragment of *CsLOB1* gene promoter region cloned in pCAMBIA1381 | This lab |
| pET41a (+) | Km^r^, IPTG-inducible expression vector | Novagen |
| pET41-CsLOB1 | Km^r^, the 714-bp full-length *CsLOB1* gene cloned in pET41a (+) | This study |
| pET41-pthA4 | Km^r^, the 3492-bp full-length *pthA4* gene cloned in pET41a (+) | This study |
| pMAL-c4x-pthA4 | Ap^r^, the 3492-bp full-length *pthA4* gene cloned in pMAL-c4x | This lab |
| pCAMBIA1300-GFP-nLUC | Km^r^, the 720-bp full-length *gfp* gene cloned in pCAMBIA1300-nLUC | This lab |
| pCAMBIA1300-cLUC-GFP | Km^r^, the 720-bp full-length *gfp* gene cloned in pCAMBIA1300-cLUC | This lab |
| pCAMBIA1300-pthA4-nLUC | Km^r^, the 3492-bp full-length *pthA4* gene cloned in pCAMBIA1300-nLUC | This study |
| pCAMBIA1300-cLUC-CsLOB1 | Km^r^, the 714-bp full-length *CsLOB1* gene cloned in pCAMBIA1300-cLUC | This study |
| CTV33 | Km^r^, Citrus tristeza virus (CTV)-based expression vector constructed in pCAMBIA1380 | El-Mohtar and Dawson, 2014 |
| CTV-2620i | Km^r^, the 488-bp *Cs9g12620* gene fragment cloned in CTV33 for gene silencing | This study |
| CTV-LOB1i | Km^r^, the 264-bp *CsLOB1* gene fragment cloned in CTV33 for gene silencing | This study |

b. List of Primers used in this study

| **Primer pair** | **Sequence (5’-3’)** | **Description or purpose** |
| --- | --- | --- |
| **Primers used for plasmid construction** | | |
| Cs9g12620.F  Cs9g12620.R | TGCTGCAGATGGCTTGGTTTCTCTCTCTT  TCGAGCTCCTATCGGAGAGGAAGAACACTAA | The 1122-bp coding sequence of *Cs9g12620* cloned in pHB vector at *Pst*I and *Sac*I sites |
| CsLOB1.F  CsLOB1.R | TGCTGCAGATGGAATGCAAACACAAAATT  TCGAGCTCTCATGTCCACAGAGGCTCC | The 714-bp coding sequence of *CsLOB1* cloned in pHB vector at *Pst*I and *Sac*I sites |
| 435-P*Cs9g12620*.F  435-P*Cs9g12620*.R | GGGGACAAGTTTGTACAAAAAAGCAGGCTTCAAGTCAACATTAGCACGAAC  GGGGACCACTTTGTACAAGAAAGCTGGGTCAAGAGAGAGAAACCAAGGCAT | A 463-bp DNA fragment of *Cs9g12620* gene promoter cloned in pGWB435-LUC with gateway |
| P*Cs9g12620*-1.R  P*Cs9g12620*-2.F | TGATTTGCTCCTATAGTCTT  AAGACTATAGGAGCAAATCAAACCAATGAAGCTTCATTTG | To obtained fragment P*Cs9g12620*-M_LB1_ and P*Cs9g12620*-M_LB1/2_  To obtained fragment P*Cs9g12620*-M_LB1_ and P*Cs9g12620*-M_LB1/2_ |
| P*Cs9g12620*-3.R | TGAGATGTGCCACTTGGTCT | To obtained fragment P*Cs9g12620*-M_LB2_ and P*Cs9g12620*-M_LB1/2_ |
| P*Cs9g12620*-4.F | AGACCAAGTGGCACATCTCATAATTTGGTGGCCCTATAAC | To obtained fragment P*Cs9g12620*-M_LB2_ and P*Cs9g12620*-M_LB1/2_ |
| 435-P*Cs9g12620*-5.R | GGGGACCACTTTGTACAAGAAAGCTGGGTCGAATGAAAACGCTCAACGGG | To construct pGWB435-P*Cs9g12620*-M |
| 435-P*Cs9g12650*.F | GGGGACAAGTTTGTACAAAAAAGCAGGCTTCGTACTTCTTGGATTT | A 450-bp DNA fragment of *Cs9g12650* gene promoter cloned in |
| 435-P*Cs9g12650*.R | GGGGACCACTTTGTACAAGAAAGCTGGGTCTTTTCTATGATGAAG | pGWB435-LUC with gateway |
| Xho-P*Cs9g12620*.F  Bam-P*Cs9g12620*.R | TGCTCGAGAAGTCAACATTAGCACGAAC  TCGGATCCAAGAGAGAGAAACCAAGCCAT | A 463-bp DNA fragment of *Cs9g12620* gene promoter cloned in pG221 at *Xho*I and *Bam*HI sites |
| Eco-P*Cs9g12620*.F  Bam-P*Cs9g12620*.R | TCGAATTCAAGTCAACATTAGCACGAAC  TCGGATCCAAGAGAGAGAAACCAAGCCAT | A 463-bp DNA fragment of *Cs9g12620* gene promoter cloned in pCAMBIA1381 at *Eco*RI and *Bam*HI sites |
| P*Cs9g12620*-A*.*F | TTCCTTTCTTTTTGTCACCCACTTTAATATATAA | To obtained fragment P*Cs9g12620*-M_A_ |
| P*Cs9g12620*-A*.*R | GTGACAAAAAGAAAGGAA |  |
| P*Cs9g12620*-C*.*F | TTCCTTTCTTTTTGTCCCCCACTTTAATATATAA | To obtained fragment P*Cs9g12620*-M_C_ |
| P*Cs9g12620*-C*.*R | GGGACAAAAAGAAAGGAA |  |
| P*Cs9g12620*-G*.*F | TTCCTTTCTTTTTGTCGCCCACTTTAATATATAA | To obtained fragment P*Cs9g12620*-M_G_ |
| P*Cs9g12620*-G*.*R | GCGACAAAAAGAAAGGAA |  |
| FL | TCCCACTTTAATATATAA | EMSA |
| FL:A | ACCCACTTTAATATATAA | EMSA |
| FL:C | CCCCACTTTAATATATAA | EMSA |
| FL:G | GCCCACTTTAATATATAA | EMSA |
| Nde-A4.1.F  Eco-A4.1.R | TTCATATGGATCCCATTCGTTCGCG  TTGAATTCGTGTGTAAACCCATGGCC | A 571-bp DNA fragment of *pthA4* gene N terminus cloned in pGBKT7 at *Nde*I and *Eco*RI |
| Eco-A4.2.F  Sal-A4.2.R | TTGAATTCGGGATGAGCAGGCACG  TTGTCGACTCACTGAGGCAATAGCTCCAT | A 561-bp DNA fragment of *pthA4* gene C terminus cloned in pGBKT7 and pET41:avrXa7 at *Eco*RI and *Sal*I sites |
| P23.F  P23.R | CACTGCAGTATTTGGTTTTACAACAACGG  GCATGCGAATTCAATTCAAACCTAGTAAATG | A 1930-bp DNA fragment amplified from CTV33 in pMD19-T Simple vector |
| End.F  End.R | TCGAATTCGCATGCTTGAAGTGGACGGAATAAG  CGATTTAAATCCCGTTTCGTCCTTTAGG | A 365-bp DNA fragment amplified from CTV33 in pMD19-T Simple vector |
| GFP.F  GFP.R | CGGAATTCTCTAGAGCGGCCGCATGGCTAGCAAAGG  CATGCATGCGGTACCCGATCGCTATTTGTAGAGCTC | A 720-bp DNA fragment amplified from CTV33 in pMD19-T Simple vector |
| Cs9g12620i.F  Cs9g12620i.R | TCGAATTCCAGCAACTCGTCCATCTC  TCGGTACCAAGAGAGGTCCACAAGTG | A 488-bp *Cs9g12620* gene fragment cloned in CTV33 vector for RNA silencing |
| LOB1i.F | TGGAATTCATGGAATGCAAACACAAAATT | A 264-bp *CsLOB1* gene fragment cloned in CTV33 vector for RNA silencing |
| LOB1i.R | TCGGTACCGCGGAGGATTTTGCAAGC |  |
| Eco-CsLOB1.F  Sal-CsLOB1.R | TGGAATTCATGGAATGCAAACACAAAATT  TCGTCGACTCATGTCCACAGAGGCTCC | A full length of *CsLOB1* gene fused in pGBKT7 and pET41a (+) Vector at *Eco*RI and *Sal*I sites |
| CsLOB1-cLUC.F | cggggcggtacccgggatccaATGGAATGCAAACACAAAATT | A full length of *CsLOB1* gene fused in pCAMBIA1300-cLUC with overlapping |
| CsLOB1-cLUC.F | cgaaagctctgcaggtcgacTCATGTCCACAGAGGCTCC |  |
| SacI-pthA4.1.F | TCGAGCTCATGGATCCCATTCGTTCGCG | A 571-bp DNA fragment of *pthA4* gene N terminus cloned in pCAMBIA1300-nLUC at *Sac*I and *Kpn*I sites |
| KpnI-pthA4.1.R | TTGGTACCGTGTGTAAACCCATGGCC |  |
| KpnI-pthA4.2.F | TTGGTACCGGGATGAGCAGGCACG | A 561-bp DNA fragment of *pthA4* gene C terminus cloned in pCAMBIA1300-nLUC at *Kpn*I and *Sal*I sites |
| SalI-pthA4.2.R | TTGTCGACCTGAGGCAATAGCTCCAT |  |
| **Primers for real time PCR analysis** | | |
| *CsActin* | CCAAGCAGCATGAAGATCAA  ATCTGCTGGAAGGTGCTGAG | 101-bp |
| *Cs9g12620* | GGTCAGTCGGGTAACCTCTCAGT  CAGAAGTTGCCTTAAAAGCCCAT | 85-bp |
| *Cs9g12650* | CTCGAGTTATTCATCCTC | 132-bp |
|  | TGATTTCCCATTTGGGAA |  |
| *CsLOB1* | AGGAACTGCCAGAATCTCAACGA  GGATTCTGGCACTTGCTTCATA | 70-bp |
| *NbEF1α* | TGGTGTCCTCAAGCCTGGTATGGTTG  ACGCTTGAGATCCTTAACCGCAACATTCTT | 155-bp |
| *gusA* | TAGAAACCCCAACCCGTGAA  TTGCCCGGCTTTCTTGTAAC | 120-bp |
| *pthA4* | TGCCCCCTCACCTGCGTTCT  CTGTGGCAGCCTCTGTATGGTGA | 131-bp |
| *pectin esterase* | TGGGTGAGTAGGGAGACGAG | 114-bp |
|  | AATCGCTTCCGCAATCGTTG |  |
| *expansin* | TCCAGCATGTTCAGGCAGAG | 115-bp |
|  | CATGGCACCGAAAGCTGTTC |  |

The 5′ end of each primer for molecular cloning contains restriction enzyme sites
